# Dynamic changes in spatial representation within the posterior parietal cortex in response to visuomotor adaptation

**DOI:** 10.1101/2021.10.18.463296

**Authors:** S Schintu, DJ Kravitz, EH Silson, CA Cunningham, EM Wassermann, S Shomstein

**Affiliations:** Behavioral Neurology Unit, National Institute of Neurological Disorders and Stroke, Bethesda, MD 20892 USA; Department of Psychological and Brain Sciences, The George Washington University, Washington, DC 20052 USA; Laboratory of Brain and Cognition, Section on Learning and Plasticity, National Institute of Mental Health, Bethesda, MD 20814 USA; Department of Psychology, School of Philosophy, Psychology and Language Sciences, The University of Edinburgh, Edinburgh, UK

**Author notes:** **Corresponding author:** Selene Schintu, Ph.D. National Institute of Neurological Disorders and Stroke, Building 10, Room 7D48, 10 Center Drive, MSC 1440, Bethesda, MD 20892-1430.

## Abstract

Recent studies used fMRI population receptive field (pRF) mapping to demonstrate that retinotopic organization extends from primary visual cortex to ventral and dorsal visual pathways by quantifying visual field maps, receptive field size, and laterality throughout multiple areas. Visuospatial representation in the posterior parietal cortex (PPC) is modulated by attentional deployment, raising the question of whether spatial representation in the PPC is dynamic and flexible and that this flexibility contributes to visuospatial learning. To answer this question, changes in spatial representation within PPC, as measured with pRF mapping, were recorded before and after visuomotor adaptation. Visuospatial input was laterally manipulated, rightward or leftward, via prism adaptation, a well-established visuomotor technique that modulates visuospatial performance. Based on existing models of prism adaptation mechanism of action, we predicted left prism adaptation to produce a right visuospatial bias via an increasing pRF size in the left parietal cortex. However, our hypothesis was agnostic as to whether right PPC will show an opposite effect given the bilateral bias to right visual field. Findings show that adaptation to left-shifting prisms increases pRF size in both PPCs, while leaving space representation in early visual cortex unchanged. This is the first evidence that prism adaptation drives a dynamic reorganization of response profiles in the PPC. Our results show that spatial representation in the PPC not only reflects changes driven by attentional deployment but dynamically changes in response to visuomotor adaptation. Furthermore, our results provide support for using prism adaptation as a tool to rehabilitate visuospatial deficits.

**SIGNIFICANCE STATEMENT:** Representation of space in the posterior parietal cortex (PPC) is reflective of attentional selection. Deploying attention to a certain spatial location modulates response profiles of a corresponding set of neurons with receptive fields in the corresponding spatial location. Here, we show that visuomotor adaptation accomplished through left-shifting prism adaptation dynamically changes the representation of space in the PPC while preserving early visual representations. This is the first evidence that visuomotor adaptation drives an instant reorganization of spatial representation in PPC. This result has implications for understanding the dynamic organization of spatial representation in the PPC and for the rehabilitation of patients with neglect syndrome, whose spatial attention is compromised.

## INTRODUCTION

The visual cortex encodes the external environment retinotopically, such that adjacent locations in space are represented by adjacent pools of neurons in the cortex (Holmes, 1918; Horton and Hoyt, 1991). This retinotopic organization extends to the dorsal and ventral visual pathways (Sereno et al., 2001; Silver et al., 2005; Swisher et al., 2007; Kravitz et al., 2013; Uyar et al., 2016), as well as to regions associated with visuospatial attention, such as the frontal eye fields (FEF) and posterior parietal cortex (PPC; Bruce and Goldberg, 1985; Schall et al., 1995; Silver and Kastner, 2009; Szczepanski et al., 2010).

Once squarely in the realm of neurophysiology, recent advances in functional magnetic resonance imaging (fMRI) enable fine-grained measures of spatial representation within the early visual cortex and PPC using population receptive field (pRF) mapping (Dumoulin and Wandell, 2008), which provides, at the voxel level, estimates of the receptive field size and location in visual space. With this technique, Sheremata & Silver (2015) showed that varying the degree of attentional engagement changes spatial representation in the PPC. This study was the first to demonstrate that spatial representation, as measured by pRF mapping in both left and right PPC, expands when attention is deployed compared to a fixation task. While this finding provides evidence for internally initiated spatial representation changes in the PPC, it is unclear whether such changes are driven only by engaging top-down attentional selection (Sheremata and Silver, 2015) or whether space representation can be influenced by changing external factors such as manipulation of the visuospatial input during visuomotor adaptation. It is unknown whether spatial representation is malleable enough to be modulated by visuomotor learning. Here, we directly probe whether spatial representation in the PPC, as measured by pRF mapping, is altered in healthy individuals in response to spatially modified visual input induced through visuomotor adaptation when a stimulus is attended vs. fixated. In a between-subject study, changes in pRFs were measured before and after the spatial input was manipulated, either leftward or rightward, with the well-established prism adaptation (PA) technique (Helmholtz, 1867; Rossetti et al., 1998). PA is a visuomotor training consisting of adaptation to displaced vision that induces directional visuospatial biases according to the direction of displacement. Left PA, by shifting visual information n-number of visual degrees from the right to the left visual field, induces a rightward visuospatial aftereffect in healthy individuals (Schintu et al., 2014, 2017; Michel, 2016). Importantly, this rightward aftereffect is similar to neglect patients’ behavior who, following right parietal lobe damage, exhibit a pathological rightward bias. On the other hand, while right PA ameliorates neglect symptoms in patients by reducing their rightward pathological bias (e.g., Rossetti et al., 1998), it generally does not have significant behavioral consequence in healthy individuals, despite inducing detectable neural changes (Crottaz-Herbette et al., 2014; Schintu et al., 2014, 2018, 2020).

We predicted that spatial representation is dynamic and, based on previous models (Pisella et al., 2006; Striemer and Danckert, 2010), expected adaptation to left-deviating prisms to produce a rightward visuospatial bias reflected by increasing representation of space in the left PPC and possibly decreasing it in the right, which however due to its effect of both visual fields would not affect behavior (Mesulam, 1981). Whereas adaptation to right-deviating prisms was expected to mirror such neural changes without altering visuospatial performance (consistent with previous behavioral literature). Since PA is believed to modulate higher-level spatial attention by acting on the PPC and given that attention enhances the neural response of the early visual areas by prioritizing regions of space (Kastner, 1998) it is possible that such pRF changes, albeit in smaller magnitude, would be also observed in early visual cortex.

The existence of rapid and dynamic changes in response of visuospatial manipulation would serve to not only constrain our current understanding of neural instantiation of attentional control but also to pave the way for promising rehabilitative techniques of compromised visuospatial processing.

## MATERIALS AND METHODS

### Participants

Forty adults with normal or corrected-to-normal vision and no history of neurological problems were recruited for the study. All participants were right-handed (Edinburgh Inventory; Oldfield, 1971), had a right dominant eye (hole-in-card test; Miles, 1930), were compensated for participation, and gave informed consent. The study was approved by the National Institutes of Health Institutional Review Board. Twenty participants were adapted to left PA (12 females; age = 26.25 ± 0.87 SEM) and the remaining twenty were adapted to right PA (13 females; age = 26.12 ± 1.05 SEM). Two participants were excluded from the right PA group, one for incomplete data collection and one because of the presence of a cyst. After pRF analysis criteria were satisfied (see pRF methods section), the final data submitted to the statistical analysis were gathered from a total of 26 participants: 16 in the left PA (10 females; age = 26.06 ± 0.9 SEM) and 10 in the right PA group (6 females; age = 24.19 ± 1.3 SEM). No difference in age was observed between the two groups [*t*(24) 1.194, *p* = 0.244].

### Procedure

The experimental procedure consisted of two general sessions, one before and after PA, and each session included behavioral testing and neuroimaging scan. A baseline measurement of visuospatial cognition was assessed with perceptual line bisection and manual line bisection tasks (see Figure 1, marked as PLB and MLB). Then, in the pre PA session participants underwent a neuroimaging scan that consisted of a resting state scan (results of which are reported in Schintu et al., 2020) and a pRF scan (focus of this report), perceptual line bisection, and manual line bisection tasks, along with straight-ahead (SA) pointing task and open-loop (OL) pointing task that were used as a proxy of the PA level (Figure1). PA then followed; one group of participants was adapted to left-deviating prisms whereas the other group was adapted to the right-deviating prisms. In the post adaptation session, the straight-ahead and open-loop pointing tasks immediately (early-post) followed PA, participants then underwent another resting state and pRF scan, which was then followed by perceptual line bisection and manual line bisection tasks, and finally (late-post) by another repeat of the straight-ahead and open-loop pointing tasks.

**Figure 1.**
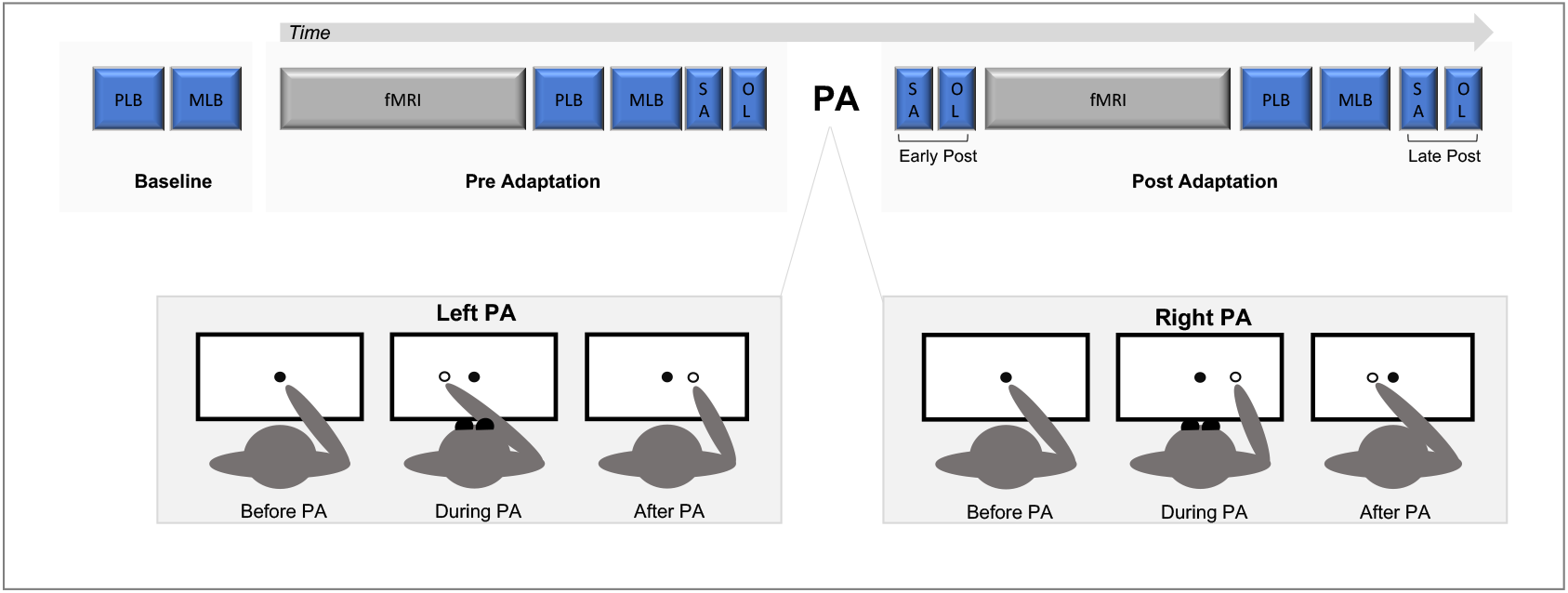
Experimental Design. PLB = perceptual line bisection task (i.e. Landmark task); MLB = manual line bisection task; fMRI = funtional magnetic resonance imaging; SA = straight-ahead pointing task; OL = open-loop pointing task; PA = prism adaptation; Pre = before prism adaptation; Post = after prism adaptation.

During behavioral assessment and PA, participants were comfortably seated in front of a horizontal wooden board with their heads supported by a chin rest. On the board, three circular targets (8 mm in diameter) were positioned at 0°, −10°, and +10° from the body midline, approximately 57 cm from participant’s nasion, and were used for PA, open-loop, and straight-ahead tasks.

### Behavioral assessment

We employed four different tasks quantifying spatial representation at the cognitive, proprioceptive, and sensorimotor level as described previously in our studies investigating visuospatial modulation following both PA and inhibitory transcranial magnetic stimulation (TMS; Schintu et al., 2020, 2021).

#### Perceptual line bisection task – The Landmark task

A modified version of the Landmark task (Milner et al., 1992) was used to measure the visuospatial shift induced by PA. The task consisted of 66 white pre-bisected lines (350 mm x ~2 mm) and were displayed on a black screen positioned 35 cm from the eyes. Lines were transected at the true center and 2, 4, 6, 8, and 10 mm toward the left and right of the true center. Each of the 11 different pre-bisected lines was presented six times in a pseudorandom order, yielding a total of 66 trials, taking approximately three minutes to complete. Each pre-bisected line was displayed for a maximum of five seconds or until a response was made and was then replaced by a black- and-white patterned mask which stayed on the screen for one second before the next pre-bisected line was displayed. Presentation software (Neurobehavioral Systems, Inc., USA) was used to generate the stimuli, record responses, and control the timing of stimulus presentation throughout the task. A screen was placed in front of the participants (35 cm from their nasion) and they were instructed to fully inspect each pre-bisected line and judge whether the mark (transector) was closer to the left or right end of the line. In this two-alternative forced-choice paradigm participants answered by pressing the pedal under their left foot if the transector was perceived as being closer to the left end of the line and by pressing the pedal under their right foot if they thought it was closer to the right end of the line. Response by pedals was chosen to limit the use of the right hand, which was used to adapt to the prisms since any feedback from that hand could contribute to de-adaptation. Participants were instructed to respond accurately and within 5 seconds. Prior to the baseline measure, at least, ten practice trials were given to ensure that participants properly understood the instructions and were confident answering with the pedals. For each participant, the percentage of ‘right’ responses was plotted as a function of the position of the transector. These data were then fitted with a sigmoid function and the value on the x-axis corresponding to the point at which the participant responded ‘right’ 50% of the time was taken as the point of subjective equality (PSE).

#### Manual line bisection task

The manual line bisection task was used to measure the visuospatial shift induced by PA (Schenkenberg et al., 1980). It consisted of a series of 10 black lines (350 mm x ~2 mm; identical in size to those used for the Landmark task) each drawn on A3 (297mm x 420mm) sheets of paper that were positioned over the computer screen which was kept at the same distance and position as for the Landmark task (35 cm from participants’ nasion). Participants were instructed to fully inspect each line and with the pen held in their right hand, draw a vertical mark where they thought the center of the line was. Once the mark had been drawn the experimenter then turned the page to reveal the next tobe-bisected line. No time limit was imposed, and participants took on average one second to place the mark on each line. For each of the ten lines, the distance between the mark placed by the participant and the true center of the line was calculated. The PSE was calculated as the average distance between the true center and the mark drawn by the participant, with marks to the right of the center coded as positive and to the left as negative.

#### Straight-ahead pointing task

The straight-ahead pointing task was used to measure the proprioceptive shift induced by PA (Rossetti et al., 1998). Participants performed six pointing movements to their perceived midline with the right index finger at a comfortable and uniform speed, while the left hand rested on the lap. Before each movement, participants received a verbal instruction to close their eyes and imagine a line splitting their body in half, project this line onto the board in front of them, point to the line while keeping their eyes closed, and then return their hand to the starting position. To ensure that participants had no visual feedback regarding either their movement or their landing position, vision of the arm and hand was occluded before movement onset by a cardboard baffle. The proprioceptive shift was measured as the average distance between the landing position and the true midline with an accuracy of +/− 0.5 cm.

#### Open-loop pointing task

The open-loop pointing task was used to measure the sensorimotor shift induced by PA (similar to Rossetti et al., 1998). Participants performed six pointing movements to the central target (0 cm) with their right index finger while resting their left hand on the lap. Participants were verbally instructed, before each pointing movement, to look at the central target, close their eyes, point to the target at a comfortable speed while keeping their eyes closed, and then return their hand to the starting position when cued by the experimenter. Similar to the straight-ahead task described above, vision of the arm and hand was occluded. The landing position of the participant’s finger was noted with a precision of +/− 0.5 cm. The sensorimotor shift was measured as the average distance between the landing position and the central target.

### Prism Adaptation (PA)

During PA, participants were fitted with prismatic goggles with either a 15° left or right visual field deviation and performed 150 verbally cued pointing movements to the right (10°) and left (−10°) targets in a pseudorandom order as called by the experimenter. Prior to each pointing movement, participants placed their right index finger on the starting position, a 1.5 cm diameter Velcro^®^ pad, placed close to the midline of their chest. Participants could not see their hand when it was in the starting position and during the first third of the pointing movement (Schintu et al., 2014). Participants were instructed to point with the index finger extended, to execute a one-shot movement at a fast but comfortable speed, and to return their hand to the starting position when instructed by the experimenter. It should be noted that while left PA produces a rightward shift in bisection judgment (McIntosh et al., 2019) it is well established that adaptation to right-deviating prisms does not produce consistent and significant behavioral modulation in healthy individuals (e.g., Schintu et al., 2017). Therefore, significant behavioral effects were expected only following left PA.

### pRF mapping

Participants were scanned on 3 Tesla Prisma (Siemens) scanner. Oblique slices were oriented on an AC-PC line. Whole-brain volumes were acquired using a 32-channel head coil, 46 slices, TR 2500ms, TE 30.0ms, voxel size 3.0 x 3.0 x 3.0mm, field of view = 192 x 138 x 192mm, 64 x 46 x 64 matrix, flip angle 70°.

### fMRI data preprocessing

All data were analyzed using the Analysis of Functional NeuroImages (AFNI) software package (http://afni.nimh.nih.gov/afni; Cox, 1996). Before pRF and statistical analyses, all images for each participant were motion corrected to the first image of the first run, detrended, and aligned to the anatomy, after removal of 2 dummy volumes to allow the magnetic field to stabilize.

The pRF mapping (Dumoulin and Wandell, 2008) experimental design was adapted from Silson et al. (2018). Stimuli were presented through a bar aperture that moved gradually through the visual field while revealing fragments of scenes (circular aperture 8.9° diameter). During each run, the bar aperture made a total of eight sweeps through the visual field (2 orientations, 4 directions). A single sweep of the visual field took 36 s and consisted of 18 separate bar positions (each 2 s). At each bar position, five images were presented (400ms each). Participants were required to maintain fixation while they performed each of two tasks. During the fixation task participants indicated via button press when the white fixation dot changed to red. Color fixation changes occurred pseudo randomly, with 4 color changes per sweep. During the attention task participants indicated via button press when a target scene fragment (yellow sunflowers field) appeared in the bar aperture. The presence of the target image occurred pseudo randomly, with four color changes per sweep. Each scanning session included six runs: three runs of the fixation task and three of the attention task. The order of the fixation or attention conditions and the hand used to answer (left or right) were counterbalanced across participants. Eye tracker recording (long-range ASL EYE-TRAC 7) was used to control fixation.

### pRF mapping analysis

The pRF mapping analysis was conducted in AFNI, using a pRF implementation for the AFNI distribution (developed by R. C. Reynolds), based broadly on previous implementations for pRF estimation (Larsson and Heeger, 2006; Dumoulin and Wandell, 2008). As in Silson et. al. (2018), the model produces asymmetric elliptical Gaussian pRF models for every possible combination of center position (x, y; 200 samples across the height and width of the screen), sigma (the size of the minor axis of the ellipse; 100 intervals overall half the screen size), aspect ratio (relative length of the major and minor axes of the ellipse; 50 even intervals from 1-5), and angle (the orientation of the major axis). Sigma (minor axis) and the major length parameter (b) were then combined to calculate the area of the ellipse (A=π*σ*b), which takes into account not only possible changes along the minor axis (σ) but also longer axis since those two are not constrained to be the same in the pRF elliptical model. This procedure results in 2 x 10^8^ possible pRFs, each of which produces a unique timeseries of response to the mapping stimulus. The model then uses both Simplex and Powell optimization algorithms to find the best time series/parameter sets (X, Y, sigma, aspect ratio) by minimizing the least-squares error of the predicted time series measured against the acquired time series in each voxel. The regions of interest (ROIs) in the visual (V1d and v,V2d and v,V3d and v) and parietal (IPS 0, 1 and 2) cortex were identified via a probabilistic atlas (Wang et al., 2015). Each subarea for a given visual or parietal ROIs were combined, the summed probability threshold was set a 70%, the minimum R^2^ was set at 0.17 (p<0.05 for a timeseries of this length) along with a minimum of 100 voxels per subject in the combined ROIs.

### Statistical analysis

Statistical analyses were performed using SPSS (IBM, Version 27.0) with alpha set at .05. All data are presented as means with the within-subjects standard error of the mean (SEM). Two-tailed paired or independent t-tests were carried out for post hoc comparisons. Effect sizes are indicated for significant effects.

## RESULTS

### Behavioral measurements

#### Perceptual line bisection – Landmark task

This task measures the contribution of perceptual rather than motor component of the visuospatial bias by asking participants to judge a series or pre-bisected lines instead of actively bisect them (Milner et al., 1992). The two pre-adaptation performances were collapsed because of the absence of a significant difference in the mean data [*t* (25) = −1.335, *p* = 0.194]. A mixed-model ANOVA with time (pre, post) as within-subjects, and group (left PA, right PA) as between-subjects, variable revealed a main effect of time [*F*(1, 24) = 7.178, *p* = 0.013, *η*^2^_*p*_= 0.230], such that both groups shifted rightward of the true center after PA (from −1.14 to 0.17 mm). No other main effect or interaction reached statistical significance [*F*s ≤ 0.311, *p*s ≥ 0.582]. Given the consistent effect of left PA on visuospatial behavior, as opposed to the right-shifting prisms which produced no significant effect, left PA group performance was submitted to a paired t-test which revealed a significant rightward shift in midline judgment from (−0.94 to 0.15 mm) following adaptation [*t*(15) = −2.142, *p* = 0.049, *Cohen’s d* = 0.535; Fig. 2 upper panel). These results show that left PA induced the anticipated rightward bias in midline judgment and thus successfully altered spatial processing.

**Figure 2.**
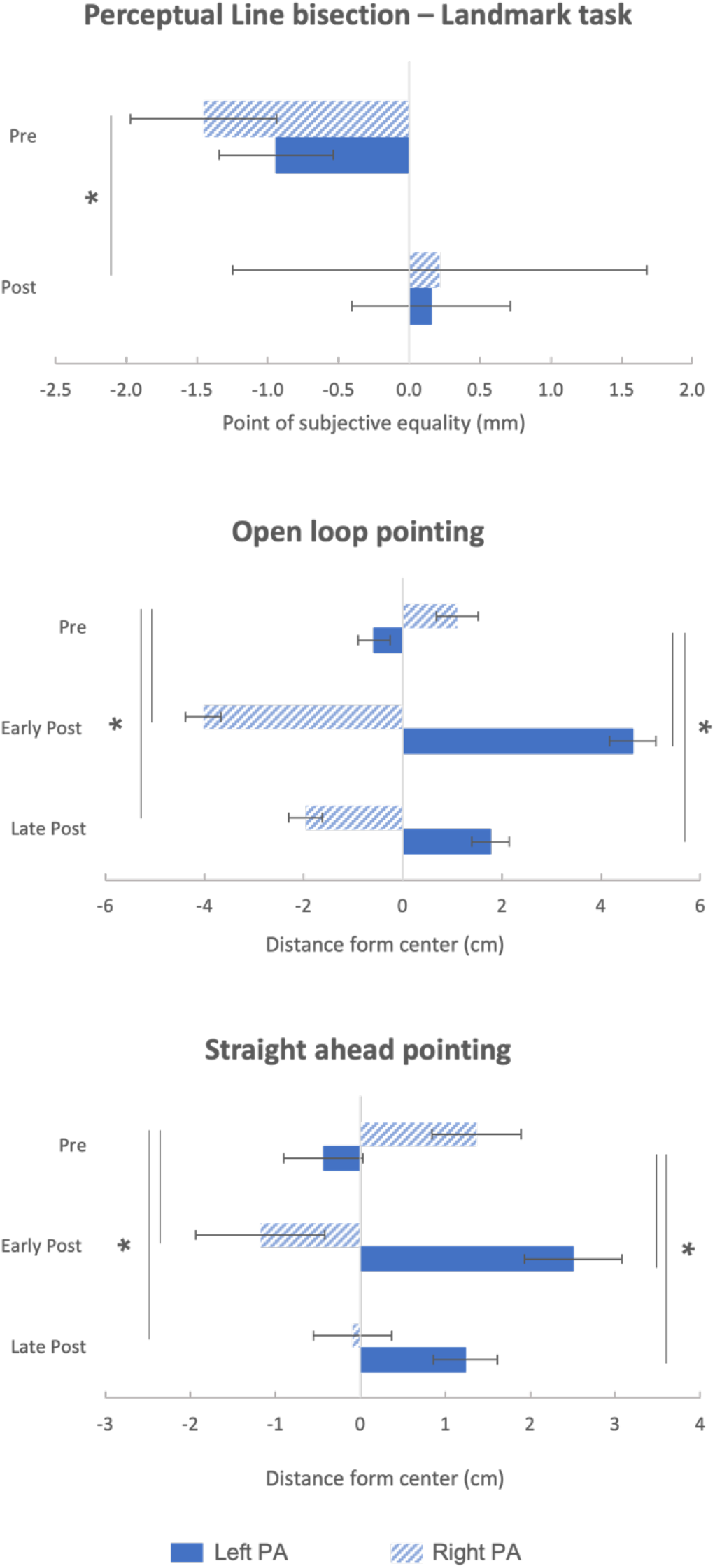
Behavioral results. Negative and positive values represent left and right of the true center (0 cm). PSE = point of subjective equality; PA = prism adaptation; Pre = before prism adaptation; Post = after prism adaptation. Error bars represent 1 standard error of the mean (SEM).

#### Manual line bisection task

This task, while perceptual in nature, quantifies the motor rather than perceptual component of the visuospatial bias (Milner et al., 1992) by measuring the difference between the perceived (manually marked) and actual center of the line. The two pre-adaptation performances were averaged, since there was no difference at baseline [*t*(25) = −1.762 *p* = 0.09]. A mixed-model ANOVA with time (pre, post) as within-subjects variable and group (left PA, right PA) as between-subjects variable revealed a trend towards a main effect of time [*F*(1, 24) = 3.607, *p* = 0.070, *η*^2^_*p*_= 0.131] such that both groups tended to shift rightward of the true center after PA (from −1.19 to −0.17 mm). No other main effect or interaction reached statistical significance [*F*s ≤ 0.367, *p*s ≥ 0.550]. These results revealed that PA, independently of the visual displacement direction, induced a weak rightward bias in midline judgments.

#### Open-loop pointing task

Sensorimotor performance before and after PA was measured by quantifying the deviation in the pointing from the landing position and the true center. A mixed-model ANOVA with time (pre, early-post, late-post) as within-subjects variable and group (left PA, right PA) as between-subjects variable revealed a significant time x group interaction [*F*(2, 48) = 226.600, *p* < 0.001, *η*^2^_*p*_= 0.904]. Planned comparisons revealed that for the left PA group the sensorimotor aftereffect from baseline (−0.58 cm) shifted rightward at early [4.63 cm; *t*(15) = −14.774, *p* < 0.001, *Cohen’s d =* −3.69] and late-post measurements [1.78 cm; *t*(15) = −8.336, *p* < 0.001, *Cohen’s d* = −2.08], and for the right PA group from baseline (1.10 cm) it shifted leftward at both early [-4.02 cm; *t*(9) = 12.310, *p* < 0.001, *Cohen’s d* = 3.89] and late-post measurements [-1.96 cm; *t*(9) = 7.921, *p* < 0.001, *Cohen’s d* = 2.50; Fig. 2 middle panel]. Importantly, this measure confirmed that PA affected behavioral visuospatial processing in the direction anticipated by left and right PA. The main effect of group [*F*(1,24) = 50.041, *p* < .001, *η*^2^*p*= .676] indicated a general rightward pointing deviation for the left PA group (1.94 cm) and leftward pointing deviation for the right PA group (−1.63 cm) whereas the main effect of time [*F*(2, 48) = 1.621, *p* = 0.208] was not significant.

To assess whether the amount of sensorimotor aftereffect differed between the two groups we compared the absolute value of the amount of change in sensorimotor performance (early- and late-post *minus* pre) between groups. The independent t-test revealed that the amount of sensorimotor adaptation between groups differed neither at early nor at late post measurements [*t*s ≤ −1.513, *p*s ≥ 0.143]. To control for the effect of the difference at baseline between the two groups [*t*(24) −3.196 *p* = 0.004 *Cohen’s d* = 1.288] we ran a mixed-model ANOVA with time (early-post, late-post) as within-subjects variable and group (left PA, right PA) as between-subjects variable and having the change in pointing (early- and late-post *minus* pre) as dependent measure and the baseline pointing as covariate. Importantly, this analysis kept revealing a significant time x group interaction [*F*(1, 23) = 96.837, *p* < 0.001, *η*^2^_*p*_= 0.808] and a non-significant time x baseline interaction [*F*(1, 23) = 0.237, *p* = 0.631], meaning that the baseline difference did not influence the changes following PA.

These results show that left PA-induced a rightward bias whereas right prims adaptation induced a leftward bias in pointing to a central visual target and that this effect has a comparable magnitude between groups, meaning that both groups were significantly and no-differently adapted until the end of the experiment.

#### Straight-ahead pointing task

Proprioceptive performance was measured by quantifying the deviation between pointing to the perceived midline and the true participants’ one. A mixed-model ANOVA with time (pre, early-post, late-post) as within-subjects variable and group (left PA, right PA) as between subject-variable revealed significant time x group interaction [*F*(2, 48) = 31.233, *p* < 0.001, *η*^2^_*p*_= 0.535]. Planned comparisons indicated that for the left PA group the proprioceptive performance from baseline (−0.43 cm) shifted rightward at both early [2.50 cm; *t*(15) = −6.338, *p* < 0.001, *Cohen’s d* = −1.58] and late (1.24 cm) post measurements [*t*(15) = −4.750, *p* < 0.001, *Cohen’s d* = −1.19], and for the right PA group from baseline (1.37 cm) it shifted leftward at both the early [-1.17 cm; *t*(9) = 4.099, *p* = 0.003, *Cohen’s d* = 1.30] and late-post adaptation measurements [-0.09 cm; *t*(9) = 2.907, *p* = 0.017, *Cohen’s d* = 0.92; Fig. 2 lower panel]. Importantly, this measure confirmed that PA affected behavioral visuospatial processing in the direction anticipated by left- and rightward shifting prisms. The main effect of time [*F*(2, 48) = 0.162, *p* = 0.851] and group [*F*(1, 24) = 2.673, *p* = 0.115] were not significant.

To assess whether the amount of proprioceptive aftereffect differed between the two groups, the absolute value of the amount of change in proprioceptive performance (early- and late-post *minus* pre) was compared between groups. An independent t-test revealed that the amount of sensorimotor adaptation between the two groups differed neither at early nor at late post [*t*s ≤ 0.162, *p*s ≥ 0.873].

To control for the possible effect of baseline difference between the two groups [*t*(24) −2.488 *p* = 0.020 *Cohen’s d* = 1.793], as for the open-loop measurement, we ran a mixed-model ANOVA with the change in pointing as dependent measure and baseline pointing as a covariate. Importantly, this analysis revealed a significant time x group interaction [*F*(1, 23) = 13.319, *p* = 0.001, *η*^2^_*p*_= 0.367] and a nosignificant time x baseline interaction [*F*(1, 23) = 3.205, *p* = 0.159].

These results show that left PA induced a rightward bias whereas right PA induced a leftward bias in pointing to the perceived midline, and that this effect was comparable in magnitude across the two groups. PA successfully altered representation of space according to the prismatic deviation.

### pRF MAPPING

Having shown that PA successfully modulated behavioral performance as measured by visuospatial, sensorimotor, and proprioceptive tasks, we examined the corresponding changes in spatial representation within the retinotopically mapped PPC (IPS 0-1-2) and sensory visual areas (V1, V2, V3) as a function of left- and right-deviating prisms. Given previous results we expected PA to alter pRF size, and more precisely it was expected to observe the largest alterations of visuospatial representation in the attention rather than fixation condition (Sheremata and Silver, 2015).

### pRF control measures

#### pRF stimuli detection

During the pRF mapping experimental run, participants performed a detection task. Performance was analyzed to ensure that the attention and fixation conditions did not differ in difficulty and that PA not affect basic visual abilities such as visual detection. The percentage of correct detection was submitted to a mixed model ANOVA with time (pre, post) and condition (attention, fixation) as within-subjects variables, and group (left PA, right PA) as a between-subjects variable. The analysis revealed solely a main effect of time [*F*(1, 24) = 7.448, *p* = 0.012, *η*^2^_*p*_= 0.237], such that both the left PA and right PA group showed a common practice effect as performance improved after adaptation (from 0.95% to 0.97%). All other main effects and interactions did not reach significance (*F*s ≤ 0.155 *p*s ≥ 0.697). This analysis confirmed that the attention and fixation condition did not differ in difficulty level, that PA did not alter visual detection abilities and that both groups improved detection with practice.

#### Eye tracking analysis

Given that attending to the stimulus could result in a larger spread of gaze position (which would increase pRF size) and since PA might affect eye position according to the direction of the lateral displacement, the standard deviation of the eye x position was submitted to a mixed-model ANOVA with time (pre, post) and condition (attention, fixation) as within-subjects variables and group (left PA, right PA) as between-subjects variable. The results for those participants for whom eye tracker data was not lost during pRF mapping (left PA = 8, right PA = 4) did not reveal any significant main effects or interactions (*F*s ≤ 1.414 all *p*s ≥ 0.262), meaning that there was no difference in accuracy in fixating the central cross across conditions and that neither left nor right PA altered it.

### pRF size

To examine changes in PPC and early visual areas, pRF size (i.e., area of the ellipse) was submitted to an omnibus mixed-model ANOVA with ROI (IPS, V1, V2, V3,) time (pre, post), condition (attention, fixation), and hemisphere (left, right) as within-subjects variables, and group (left PA, right PA) as between-subjects variables. This analysis revealed a significant 5-way interaction ROI x time x condition x hemisphere x group [*F* (3, 72) = 3.238, *p* = 0.027, *η*^2^_*p*_= 0.119], along with an ROI x condition x hemisphere x group [*F*(3, 72) = 3.066, *p* = 0.033, *η*^2^_*p*_= 0.113], ROI x time x condition x group [*F*(3, 72) = 4.373, *p* = 0.007, *η*^2^_*p*_= 0.154], ROI x time x group [*F*(3, 72) = 3.204, *p* = 0.028, *η*^2^_*p*_= 0.118], ROI x condition x hemisphere [*F*(3, 72) = 4.133 *p* = 0.009, *η*^2^_*p*_= 0.147], and ROI x condition [*F*(3, 72) =3.022 *p* = 0.35, *η*^2^_*p*_= 0.112] interaction. There were also a significant main effect of ROI [*F*(3, 72) = 5.943, *p* = 0.001, *η*^2^_*p*_= 0.198], time [*F*(1, 24) = 6.830 *p* = 0.015, *η*^2^_*p*_= 0.222], condition [*F*(1, 24) = 22.369, *p* < 0.001, *η*^2^_*p*_= 0.482], and group [*F*(1, 24) = 5.254 *p* = 0.031, *η*^2^_*p*_= 0.180]. All other main effects and interactions were not significant (*F*s ≤ 2.201 *p*s ≥ 0.095).

To unpack the 5-way interaction, the corresponding hubs in the visuospatial attention network (Szczepanski et al., 2010) were separated for further statistical analyses. Namely, parietal and early visual cortex ROIs were analyzed independently.

#### Parietal cortex

pRF size data were submitted to a mixed-model ANOVA with time (pre, post), condition (attention, fixation) and hemisphere (left, right) as within-subjects variables, and group (left PA, right PA) as between-subjects variable. The analysis revealed a significant time x condition x group interaction [*F*(1, 24) = 5.659, *p* = 0.026, *η*^2^_*p*_= 0.191], follow-up planned comparisons of the 3-way interaction revealed a modulation of the pRF size for the attention condition, which bilaterally increased (from 4.04° to 5.36°) following left PA [*t*(15) −2.202, *p* = 0.044, *Cohen’s d* = −0.551] meaning that left PA increased pRF size in both left and right PPCs (Fig. 3). The analysis revealed also a main effect of condition [*F*(1, 24) = 13.819, *p* = 0.001, *η*^2^_*p*_= 0.365] such that pRF size was larger in the attention (4.73°) as compared to fixation (3.41°) condition, replicating earlier findings (Sheremata and Silver, 2015). All other main effects and interactions were not significant (*F*s ≤ 3.986 *p*s ≥ 0.057). Overall, this analysis revealed that left PA bilaterally increased pRF size when attention was deployed.

**Figure 3.**
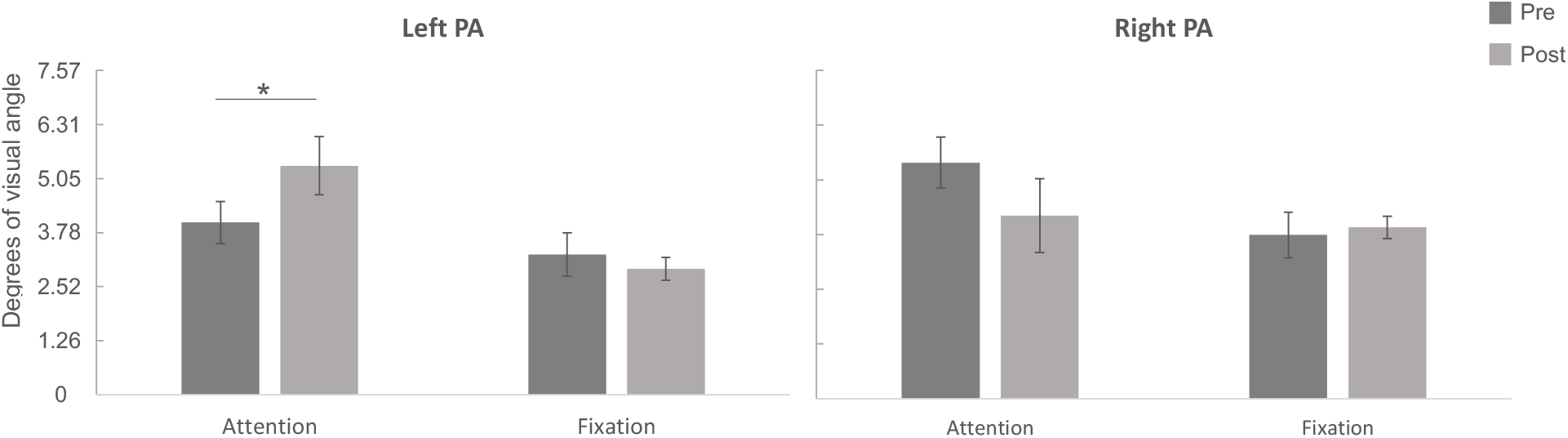
Parietal cortex. pRF computed size expressed in degrees of visual angle. PA = prism adaptation; Pre = before prism adaptation; Post = after prism adaptation. Error bars represent 1 SEM.

All other comparisons controlling for possible baseline condition differences both between groups (*t*s ≤ −1.435 *p*s ≥ 0.234) and within groups (*t*s ≤ 1.677 *p*s ≥ 0.114) did not reach statistical significance.

**Figure 4.**
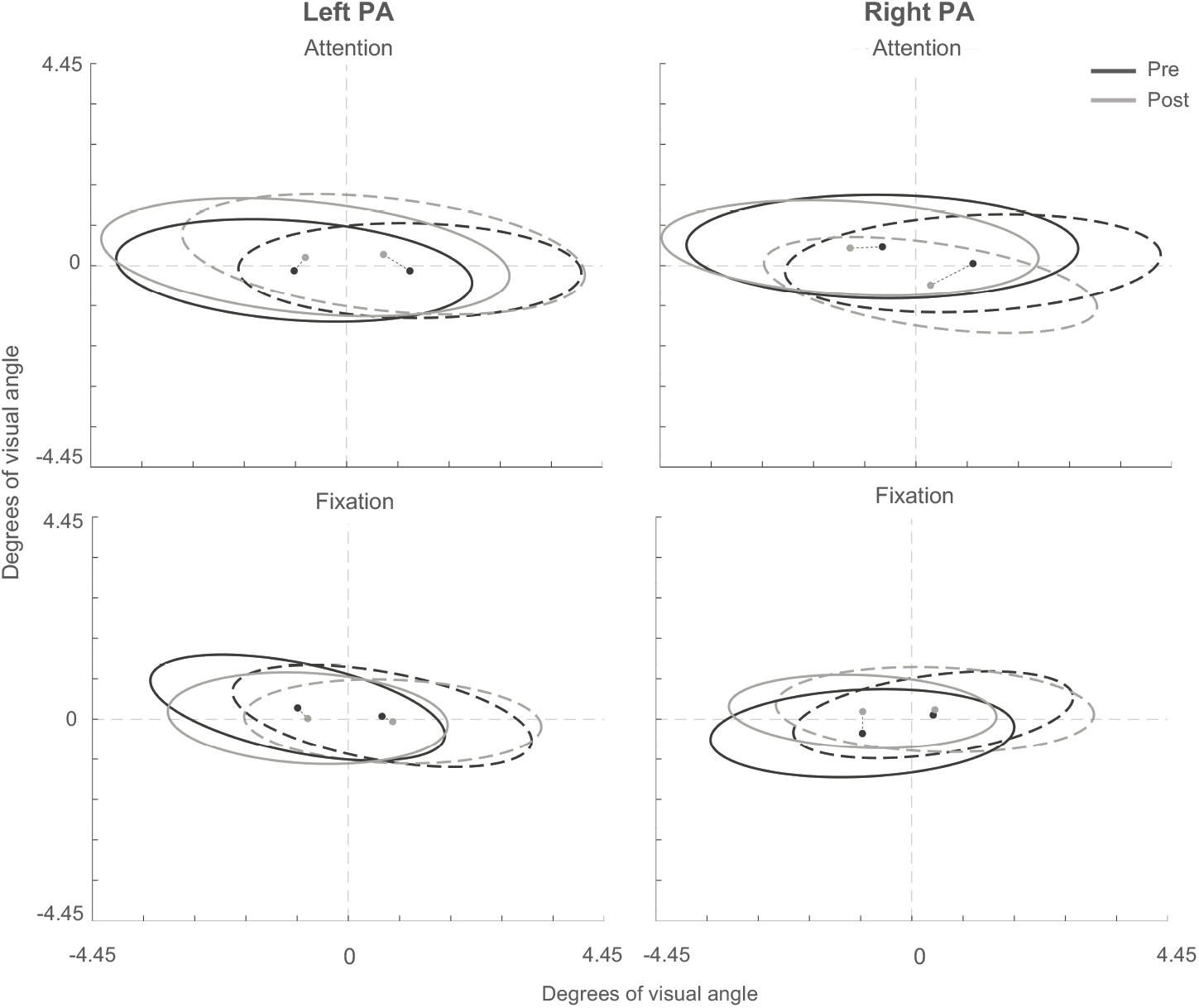
Parietal cortex. Elliptic representation of computed pRF size for each group and condition. Solid line = right hemisphere (left visual field); Dotted line = left hemisphere (right visual field). PA = prism adaptation.

#### Early visual cortex

pRF size data were submitted to a mixed-model ANOVA with ROI (V1, V2, and V3), time (pre, post), condition (attention, fixation), hemisphere (left, right) as within-subjects variables, and group (left PA, right PA) as a between-subjects variable. The analysis showed an ROI x time x condition x hemisphere interaction [*F*(2, 48) = 6.817, *p* = 0.002, *η*^2^_*p*_= 0.221] and post-hoc planned comparisons revealed that independently of the direction of the visuospatial manipulation pRF size increased over time (with practice) in V1 [left hemisphere *t*(25) −2.235, *p* = 0.035, *Cohen’s d* = −0.438], V2 [left *t*(25) −2.850, *p* = 0.009, *Cohen’s d* = −0.559 and right hemisphere *t*(25) −2.156, *p* = 0.041, *Cohen’s d* = −0.423] and V3 [right hemisphere *t*(25) −2.396, *p* = 0.024, *Cohen’s d* = −0.470]. There was also a ROI x time x group interaction [*F*(2, 48) = 4.104, *p* = 0.023, *η*^2^_*p*_= 0.146] such that solely for the left PA group V2 pRFs increased from pre (2.77°) to post (3.28°) adaptation [t(15) = −2.222, *p* = 0.042, *Cohen’s d* = −0.556; others *t*s ≤ −2.027 *p*s ≥ 0.073] meaning that, overall, pRF size had the general tendency to increase bilaterally following left PA independently of the condition. There was also a condition x hemisphere x group interaction [*F*(1, 24) = 4.340, *p* = 0.048, *η*^2^_*p*_= 0.153] such that, independently of the visuospatial manipulation and across ROIs, the right PA group showed larger pRFs in the attention condition (4.16°) than the fixation condition (3.03°) in the left hemisphere [t(9) = 3.779, *p* = 0.004, *Cohen’s d* = 1.195] along with a marginal difference between the attention condition in the right hemisphere (3.91°) and the fixation condition in the left hemisphere (3.03°) [t(9) = 2.232, *p* = 0.053, *Cohen’s d* = 0.706], this interaction also revealed that the right PA group had larger pRFs than the left PA group for the attention condition in both the right (left PA = 2.96° right PA = 3.91°; t(24) = −2.179; *p* = 0.039; *Cohen’s d* = −0.878) and left (left PA = 2.90° right PA = 4.16°; t(24) = −2.724; *p* = 0.012; *Cohen’s d* = −1.10) hemispheres (others *t*s ≤ −1.979 *p*s ≥ 0.059).

The analysis also showed a main effect of time [*F*(1, 24) = 8.836, *p* = 0.015, *η*^2^_*p*_= 0.222] such that overall pRF size increased from pre (2.84°) to post (3.41°) adaptation, a main effect condition [*F*(1, 24) = 10.728, *p* = 0.003, *η*^2^_*p*_= 0.309] such that overall pRFs were larger in the attention (3.34°) than fixation (2.90°) condition. Finally a main effect of group [*F*(1, 24) = 5.632, *p* = 0.026, *η*^2^_*p*_= 0.190] was also found, such that pRFs were overall larger in the right PA (3.66°) than left PA (2.77°) group. All other comparisons did not reach significance (Fs ≤ 3.593 *p*s ≥ 0.070; Fig.

Overall, results from the early visual cortex show that, unlike the parietal cortex, early sensory V1-V3 regions do not show any changes in pRF size exclusively driven by altering spatial input by PA. Rather, consistent with behavioral data, we observed a general tendency of pRFs to increase size over time, likely reflecting practice effects in change detection task (see *pRF control measure* section above).

### pRF size separated by x-center position, y-center position, and angle

#### pRF x-center position

To examine changes in PPC and primary visual areas pRF center position along the x axis, the x-center position parameter was submitted to an omnibus mixed-model ANOVA with ROI (IPS, V1, V2, V3,) time (pre, post), condition (attention, fixation) and hemisphere (left, right) as within-subjects variables, and group (left PA, right PA) as between-subjects variables. The analysis revealed a significant ROI x time x hemisphere interaction [*F*(3, 72) = 3.045, *p* = 0.034, *η*^2^_*p*_= 0.113] such that, independently of whether it was the fixation or attention condition or whether participants were adapted to left or right prism, solely in V2 the x-center position shifted further leftward from pre (−0.34au) to post (−0.38au) in the right hemisphere [*t*(25) 3.263, *p* = 0.003, *Cohen’s d* = 0.640] and further rightward in the left hemisphere [from 1.78° to 1.91°; *t*(25) −2.329, *p* = 0.028, *Cohen’s d* = −0.457], all others (*t*s ≤ 1.676, *p*s ≥ 0.106).

The ROI x time x condition interaction was also significant [*F*(3, 72) = 2.725, *p* = 0.050, *η*^2^_*p*_= 0.102] such that in IPS at post, the x-center position marginally differed between the attention (−0.31°) and fixation (−0.04°) condition [*t*(25) −1.983,*p* = 0.058, *Cohen’s d* = −0.389] independent of the direction of the visuospatial modulation. There was also a significant ROI x hemisphere interaction [*F*(3, 72) = 92.683, *p* < 0.001, *η*^2^_*p*_= 0.794] such that in the left hemisphere the x-position differed across all ROIs (*t*s ? −2.168, *p*s ≤ 0.040) except for V1 versus V2 [*t* (25) = −0.707, *p* = 0.486], and in the right hemisphere the x-position differed across all ROIs (*t*s ≥ 2.173, *p*s ≤ 0.049) except for V1 versus V2 [*t* (25) = −0.101, *p* = 0.920], and a condition x hemisphere interaction [*F*(1, 24) = 14.793, *p* = 0.001, *η*^2^_*p*_= 0.381] such that the x-position differed between and within condition across all ROIs (*t*s ≥ 2.476, *p*s ≤ 0.020). The analysis revealed also a main effect of hemisphere [*F*(1, 24) = 3455.77, *p* < 0.001, *η*^2^_*p*_= 0.993] such that the x-position in the right hemisphere was negative (−1.47°), thus coding the left side of the visual field and positive in the left hemisphere (1.6°), thus coding for the right side of the visual field, and a main effect of ROI [*F*(3, 72) = 5.800, *p* = 0.001, *η*^2^_*p*_= 0.195] such that the x-position of each V areas differed from IPS (*t*s ≥ 2.537, *p*s ≤ 0.018) but not difference was found within V1-V3 (*t*s ≤ −0.782, *p*s ≥ 0.442). All others main effect and interaction were not significant (*t*s ≤ 1.914, *p*s ≥ 0.135).

Overall, the x-center position results show no changes in such parameters due to the direction of the visuomotor adaptation, but rather a change over time restricted to V2 area that was also independent of task condition.

#### pRF y-center position

To examine changes in PPC and primary visual areas pRF center position along the y axis, the y-center position parameter was submitted to an omnibus mixed-model ANOVA with ROI (IPS, V1, V2, V3,) time (pre, post), condition (attention, fixation) and hemisphere (left, right) as within-subjects variables, and group (left PA, right PA) as between-subjects variables. The analysis revealed a significant ROI x time x condition x group interaction [*F*(3, 72) = 4.393, *p* = 0.007, *η*^2^_*p*_= 0.155], planned comparison revealed that solely for V1 and following left PA the y-position increased/moved further up in the visual field in both the attention [from 0.24° to 0.40°; *t*(15) = −2.270, *p* = 0.038, *Cohen’s d* = −0.568] and fixation [from 0.09° to 0.22°; *t*(15) = −2.789, *p* < 0.014, *Cohen’s d* = − 0.697] condition (all others *t*s ≤ 1.513 *p*s ≥ 0.151). The analysis also showed an ROI x hemisphere interaction [*F*(3, 72) = 3.133, *p* = 0.031, *η*^2^_*p*_= 0.115] driven by an overall left (−0.53°) versus right (−0.27°) hemisphere difference solely for V2 [*t*(25) = −3.690, *p* < 0.001, *Cohen’s d* = −0.724], and a main effect of ROI [*F*(3, 72) = 62.843, *p* < 0.001, *η*^2^_*p*_= 0.724] such that the y-position differed across all ROIs (*t*s ≥ 4.437, *p*s ≤ 0.001) except between IPS and V1 [*t*(25) – 1.639, *p*= 0.114]. All others main effect and interaction did not reach significance (*F*s ≤ 2.752, *p*s ≥ 0.110).

Overall, this analysis showed that solely in V1 following adaptation to left-shifting prims the y-center position bilaterally increased/moved further up in the visual field over time, independently of whether pRFs were quantified during the attention of fixation task. We do not have a further interpretation of this result as we had no a-priory hypothesis regarding a shift in y-center position. Further investigations are warranted to examine the putative shift in y-center in V1.

#### pRF angle

To examine changes in PPC and primary visual areas pRF the angle parameter, for which 0° represents the horizontal axis with positive angles toward the upper vertical meridian and negative angles toward the lower vertical meridian, was also submitted to an omnibus mixed-model ANOVA with ROI (IPS, V1, V2, V3,) time (pre, post), condition (attention, fixation) and hemisphere (left, right) as within-subjects variables, and group (left PA, right PA) as between-subjects variables. This analysis showed an ROI x group [*F*(3, 72) = 4.512, *p* = 0.006, *η*^2^_*p*_ = 0.158] interaction such that solely in IPS, independently of time, condition and hemisphere, the angle parameter was negatively biased for left PA (−10.31°) and positively biased for right PA (1.72°) group [*t*(24) = − 2.340, *p* = 0.028, *Cohen’s d* = − 0.914]. There was also an ROI x hemisphere interaction [*F*(3, 72) = 38.616, *p* < 0.001, *η*^2^_*p*_ = 0.617] such that for all visual areas the angle was positively biased for the left hemisphere and negatively for the right hemisphere (*t*s ≥ 3.657 *p*s ≤ 0.001), but no difference was found in IPS [*t*(25) = 0.013, *p* = 0.990]. A main effect of hemisphere was also found [*F*(1, 24) = 46.780, *p* < 0.001, *η*^2^_*p*_ = 0.661] such that overall the left hemisphere had a positively biased angle (3.44°) and the right hemisphere a negatively biased one (−10.89°). All other main effects and interactions did not reach significance (*F*s ≤ 1.629, *p*s ≥ 0.190). Overall, this analysis revealed no changes in angle parameter due to the visuomotor manipulation.

### pRF Sigma in PPC

Given that most prior work on pRF in PPC was based on the well-established 2-D gaussian modeling (e.g., Dumoulin and Wandell, 2008; Silson et al., 2015), as opposed to the most recent elliptical one (Silson et al., 2018), we also examined changes in PPC considering only the sigma parameter (i.e., the shorter axis of the ellipse).

We submitted the pRF sigma parameter to a mixed-model ANOVAs with time (pre, post), condition (attention, fixation), and hemisphere (left, right) as within-subjects variables, and group (left PA, right PA) as between-subjects variable. The analysis revealed a significant interaction time x condition x hemisphere x group interaction [*F*(1, 24) = 4.240, *p* = 0.05, *η*^2^_*p*_= 0.150], follow-up planned comparisons showed a significant increase (from 1.07° to 1.25°) for the attention condition in the left hemisphere following left PA [*t*(15) −2.241 *p* = 0.041 *Cohen’s d* = 0.069], and a marginally significant decrease (from 1.25° to 0.98°) after right PA [*t*(15) 2.135 *p* = 0.062 *Cohen’s d* = 0.062] (Fig. 6a). This analysis also revealed a main effect of condition [*F*(1, 24) = 9.228, *p* = 0.006, *η*^2^_*p*_ = 0.278] such that overall the pRF sigma parameter was larger in the attention (1.16°) as compared to fixation (1.02°) condition, replicating earlier findings (Sheremata and Silver, 2015). All other main effects and interactions were not significant (*F*s ≤ 4.075 *p*s ≥ 0.055). All other comparisons controlling for possible baseline condition differences both between groups (*t*s ≤ −233 *p*s ≥ 0.229) and within groups (*t*s ≤ 1.723 *p*s ≥ 0.105) did not reach statistical significance. These results show that spatial representation in PPC dynamically changes in response to alteration of the spatial input. Interestingly, the sigma parameter results, despite largely overlapping with the size parameter ones, seem more sensitive to the visuomotor manipulation induced effect. Further experiments utilizing more in-depth pRF methods (i.e., symmetric vs. asymmetric pRF estimates) are warranted to adjudicate between different pRF parameters and models (Dumoulin and Wandell, 2008; Silson et al., 2018).

**Figure 5.**
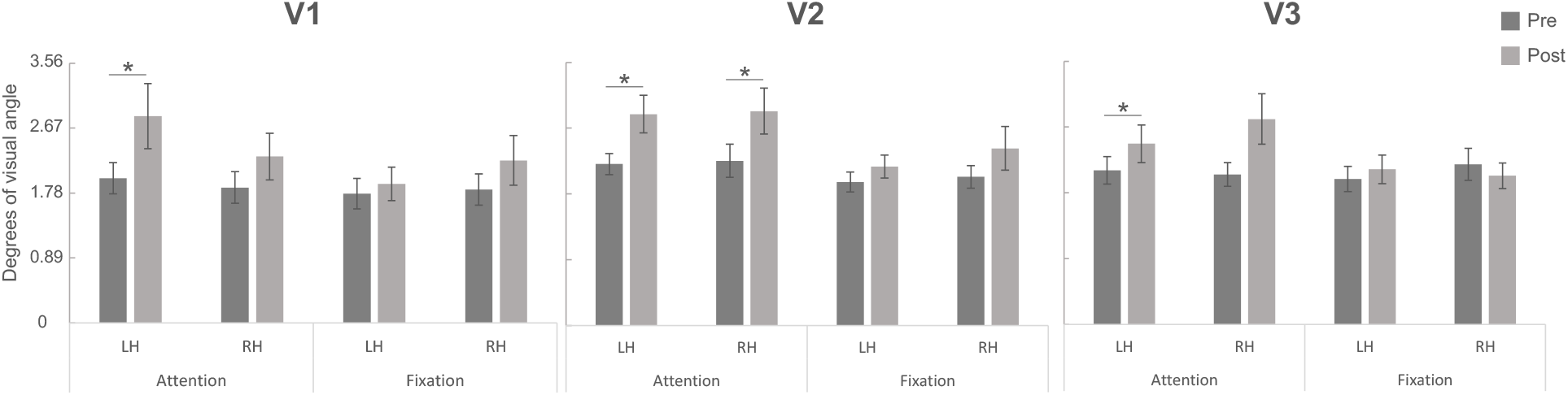
Early visual cortex. pRF calculated size expressed in degrees of visual angle. Pre = before prism adaptation; Post = after prismatic adaptation; LH = left hemisphere; RH = right hemisphere. Error bars represent 1 SEM.

**Figure 6.**
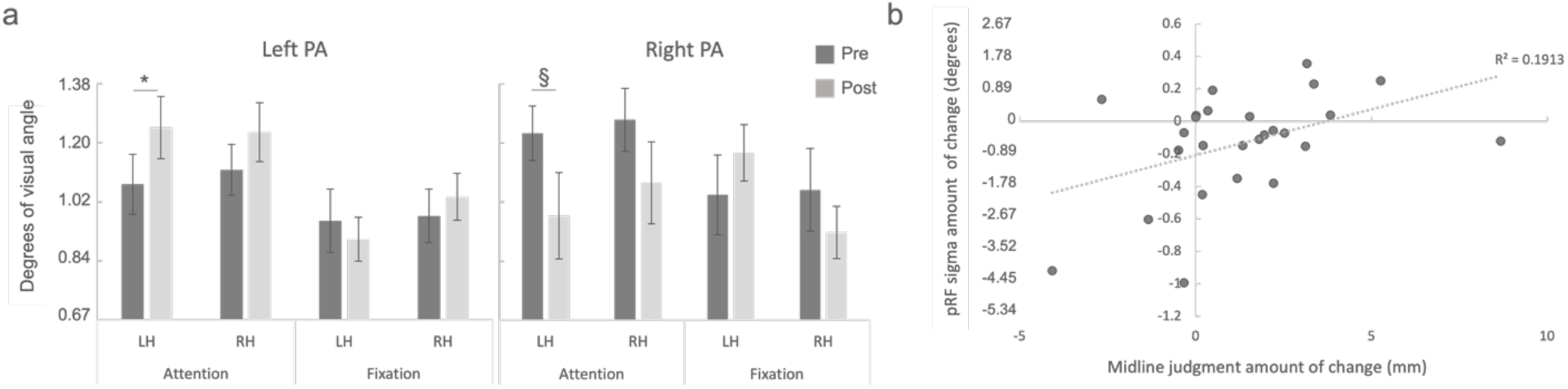
**a) Parietal cortex.** pRF sigma expressed in degrees of visual angle. PA = prism adaptation; Pre = before PA; Post = after PA; LH = left hemisphere; RH = right hemisphere. Error bars represent 1 SEM. 8 = p< 0.05; §= p < 0.06. **b) Brain behavior correlation.** Pearson correlation between the change in pRF size in the left hemisphere for the attention condition and the shift in midline judgment at the Landmark task.

### Correlation between neural and behavioral changes

To investigate the possible relationship between neural and behavioral changes following PA we computed a Pearson correlation between the change in both pRF sigma and size parameters and the shift in line bisection judgment as measure by the Landmark task. The analysis revealed that, across both groups, pRF sigma changes in the left hemisphere for the attention condition predicted the shift in perceptual midline judgment (*r* = 0.44, *p* = 0.025), meaning that the increase in pRF size in the left PPC following PA is predicted by a rightward shift as measured by the Landmark task (Fig. 6b). No significant correlation was found instead when the left PA group bilateral calculated size amount of change was correlated to midline bisection judgment amount of change (*r* = −0.218, *p* = 0.418).

## DISCUSSION

The aim of the study was to investigate whether the representation of visual space in the PPC is modulated in response to manipulation of the visual input induced by visuomotor adaptation. Using behavioral methods, it was first confirmed that PA altered visuospatial performance. Then, changes in the PPC and early visual cortex were quantified by the asymmetrical pRF model before and after left or right PA. Based on the behavioral modulation produced by left PA in healthy individuals (Colent et al., 2000; Schintu et al., 2014) and previous models (Pisella et al., 2006; Striemer and Danckert, 2010), we anticipated left PA to produce a rightward visuospatial bias by mainly increasing the representation of space in the left PPC, since any change in the right PPC would impact both visual fields (Mesulam, 1981). Opposite neural changes were expected following right PA despite the absence of corresponding behavioral ones (Pisella et al., 2006; Striemer and Danckert, 2010). In agreement with the prediction, the results showed that left PA increases pRF size in both PPCs when attention is deployed, therefore increasing the representation of visual space in the left hemisphere, and thus possibly allocating more attention to the right side of space.

### Behavioral changes

The effect of PA on visuospatial behavior, as expected, revealed that the group adapted to left-shifting prism exhibited a significant rightward bias as measured by the Landmark task; the subjective midline judgment significantly shifted rightward as compared to the baseline measurement (Fig. 2). The absence of a behavioral modulation following adaptation to right-shifting prisms is not surprising given the established observation that it fails to produce significant changes in behavior (Schintu et al., 2017) while however producing neural changes (Crottaz-Herbette et al., 2014; Schintu et al., 2020). Finding in the literature reporting the absence of behavioral modulation in healthy individuals following adaptation to right-shifting prisms has not been discussed but rather accepted as a fact, so much that right PA has become the control condition for the left one. The absence of a significant modulation at the manual line bisection task in both groups was not surprising since this is an ideal screening tool for neglect’s symptoms but its sensibility to detect visuospatial alteration in healthy is lower than the Landmark task (McIntosh et al., 2019). When considering the open-loop and straight-ahead pointing tasks, proxy of PA level, the groups’ sensorimotor and proprioceptive aftereffect did not differ (Fig. 2), ensuring that any between groups difference, at both behavioral and neural level, could not be ascribed to a difference in the adaptation strength produced by the visual shift direction. Both groups were also significantly adapted until the end of the experiment as evidenced by the two pointing tasks’ performance, ensuring that while the behavioral and neural data were acquired participants were in a state of adaption (Fig. 2).

### Neural changes

Changes in spatial representation were tracked in the PPC as this is the region that has spatial topography (Silver and Kastner, 2009) and is sensitive to changes in spatial representation following attentional modulation (Sheremata and Silver, 2015). Importantly, here, we further restricted our analysis to the three purely visuospatial subregions, IPS 0-1-2, that have been shown to affect visuospatial performance when perturbed by TMS (Szczepanski and Kastner, 2013).

At the center of our investigation was the question of whether spatial representation in the PPC, as measured by pRF, would show dynamic updating to reflect visuospatial adaptation. Consistent with the prediction, we observed bilateral (left and right hemisphere) changes in pRFs sizes following left PA. Supporting the hypothesis that PA particularly affects attentional allocation, changes in pRF sizes were observed exclusively in the attention, not fixation, condition. Namely, pRF size for the attention condition increased after adaptation to left-shifting prims, meaning that, in agreement with previous findings (Sheremata and Silver, 2015), pRF size was modulated solely when attention was deployed. Importantly, the pRF modulation in the attention condition cannot be attributed to the different task difficulty across conditions, as both groups improved their performance after adaptation independently of condition and direction of the visual displacement. Such pRF modulation cannot be explained either by eye movement since the variability of the x-position, measured via eye tracker during the experimental sessions, did not differ across time, conditions, and groups.

Unlike the PPC, early sensory V1-V3 regions did not show any pRFs size change following visuomotor adaptation. As shown in Fig. 5, pRF size increased over time independently of the direction of the visuospatial manipulation, likely reflecting a practice effect observed in the detection task, for which participants’ detection performance of pRF stimuli during the mapping improved over time independently of the PA direction. On one hand, if the changes induced by the visuomotor manipulation are attentional in nature, pRF changes following left PA would be expected in early visual areas given that attention enhances the neural response of the early visual areas by prioritizing regions of space (Kastner, 1998). On the other hand, if attentional and visuomotor adaptation would have been observed in early sensory areas our results would be open for an interpretation that it is visual changes, not attentional ones, driven by sensorimotor adaption are the ones responsible for re-organization in the PPS. Thus, the presence of pRF modulation following left PA in the PPC and its absence in early visual areas provides insight into the possible mechanism responsible for such spatial representation changes following manipulation of the visual input. The fact that changes in the PPC do not affect visual areas as previously shown (Kastner, 1998) allows us to speculate that PA-induced changes are at a higher rather than lower level of the neural processing hierarchy and that PA mechanism of action is not purely attentional but it can rather depend on local changes of spatial representation stemming from visuomotor adaption. One has to be careful, however, as not to overinterpret null results. Left PA was also found to modulate the y-center position but solely in V1 and independently of whether pRFs were quantified during the attention or fixation task. This result is unexpected and further investigations are warranted to examine the putative y-center shift in V1.

Our findings show that spatial modulation of visual input significantly altered space representation in both PPCs (Fig. 3). This can be apparently at odds with PA models initially proposed (Pisella et al., 2006; Striemer and Danckert, 2010) that, grounding on the hemispheric rivalry (Kinsbourne, 1977), hypothesized PA to modulate visuospatial cognition by inhibiting the PPC contralateral to the sensory displacement (i.e., left PA - right PPC) and in turn increase activation of the PPC ipsilateral to the displacement (i.e., left PA - left PPC). It is however in partial agreement with recent work from our laboratory along with previous studies showing neural bilateral changes in healthy individuals (Schintu et al., 2020) and neglect patients (Saj et al., 2013) following PA. Such bilateral modulation of the PPC, instead of a push-pull pattern, calls for a revision of the PA mechanism of action models. Our results are consistent with Crottaz-Herbette et al., (2014) conclusion that PA modulates spatial representation in the PPC; however, their findings show an increase of BOLD signal after right prism in the left inferior parietal lobule (IPL) while subjects were performing a visual search task. It is possible that different i) techniques employed (pRF versus event-related fMRI) and ii) level of attentional deployment required, along with iii) brain locations analyzed (IPS 0-1-2 versus IPL) may account for this discrepancy. The same reasons could explain the difference between the local pRF modulation reported here and the more widespread change in connectivity measured at rest (Schintu et al., 2020), which is, nonetheless calling further investigation to link local and global changes within the PPC in the context of spatial representation. Finally, when pRF changes in the PPC were also investigated by analyzing the sigma parameter, instead of the calculated size parameter, significant changes following left PA were restricted to the left hemisphere, and for the same hemisphere those following right PA were close to reaching significance (Fig. 6). Moreover, pRF changes, as measured by sigma, correlated with the PA-induced visuospatial bias. The sigma-based results are mostly compatible with the calculated size measure, as we can visually appreciate (Fig. 3 and 6) that the direction of change for the attention condition is bilateral (increase for left PA and decrease for right PA). However, the sigma parameter appears to show more sensitivity to the visuomotor adaptation induced modulation than the calculated size one, possibly meaning that most of the pRF modulation happens along the ellipse shorter axis.

In conclusion, here we present the first evidence of the dynamic nature of spatial representation in the PPC following visuomotor adaptation as measured with the pRF method. These results have important implications for understanding the neural instantiation of representation of space in the PPC and offer additional evidence for the usefulness of PA as a rehabilitation tool for neurological disorders exhibiting deficits in spatial allocation, such as neglect syndrome.

## ACKNOWLEDGMENT

This work was supported by the National Institutes of Health Ruth L. Kirschstein National Research Service Award (to Schintu), the Clinical Neurosciences Program of the National Institute of Neurological Disorders and Stroke (1ZIANS002977-20 to Schintu and Wassermann), and the National Science Foundation (BCS-1921415 to Shomstein, and BCS-2022572 to Shomstein & Kravitz).

